# Hidden state dynamics reveal the prolonged inactive state across the adult lifespan

**DOI:** 10.1101/2020.01.27.920652

**Authors:** Keyu Chen, Ruidi Wang, Dong-Qiang Liu

**Author notes:** **Correspondence:** Dong-Qiang Liu, Ph.D.

## Abstract

Adult lifespan is accompanied by functional reorganization of brain networks, but the dynamic patterns behind this reorganization remain largely unclear. This study focuses on modelling the dynamic process of spontaneous activity of large-scale networks using hidden Markov model (HMM), and investigates how it changes with age. The HMM with 12 hidden states was applied to temporally concatenated resting state fMRI data from two dataset of 176 / 170 subjects (aged 20-80 years), and each hidden state was characterized by distinct activation patterns of 17 brain networks. Results showed that (a) For both datasets, the elder tended to spend less time on and had less transitions to states showing antagonistic activity between various pairs of networks including default mode network, cognitive control and salience/ventral attention networks. (b) For both datasets, the elder were probable to spend more time on, have less transitions from and have more transitions to an ‘baseline’ state with only moderate-level activation of all networks, the time spent on this state also showed an U-shaped lifespan trajectory. (c) For both datasets, HMM exhibited higher specificity and reproducibility in uncovering the age effects compared with temporal clustering method, especially for age effects in transition probability. (d) These results demonstrate the age-correlated decrease of the anti-correlation between various networks, and further validate the prediction of Naik et al. (2017) that the existence of a particular network state with lower transition probability and higher fractional occupancy in old cohort, which may reflect the shift of the dynamical working point across the adult lifespan.

## 1. Introduction

The human brain undergoes structural and functional changes across the whole lifespan. Apart from the local changes, increasing efforts have been devoted to the alterations in the interaction among regional brain activity also underlie the age-related cognitive decline (O’Sullivan et al. 2001; Ferreira and Busatto 2013; Dennis and Thompson 2014; Damoiseaux et al. 2017;Zuo et al., 2017). Resting-state fMRI (rs-fMRI) is one of the most popular techniques to investigate the functional integration between large-scale networks (Biswal et al., 1995; Grecius et al. 2003; Fox and Raichle, 2007). A multitude of studies have unveiled the age-related changes in functional networks (Andrews-Hanna et al., 2007; Damoiseaux et al., 2008; Biswal et al., 2010; Tomasi and Volkow, 2012; Cao et al., 2014; Chan et al. 2014; Betzel et al. 2014; Yang et al. 2014; Geerligs et al., 2015) via static functional connectivity (sFC). Overall, thesel studies converge on the findings of decreased within- and increased between-network connectivity in the older, which may indicate decreased functional specificity or “dedifferentiation” (Cabeza 2002; Li et al. 2006; Koen et al. 2019; Edde et al. 2020) of the information processing of the human brain.

In spite of the progress, recent dynamic functional connectivity (dFC) studies have suggested that the resting brain activity is highly dynamic (Chang and Glover, 2011; Hutchison et al. 2013; Allen et al. 2014), and could be clustered into discrete and recurring ‘brain states’. Statistics like dwell time (the time spent visiting each state) and transition probability (the frequency of transition between a pair of states) were also introduced to measure state-specific network dynamics in some brain disorders (Damaraju et al. 2014; Rabany et al. 2019; Kim et al. 2017; Zhuang et al. 2018;De Vos et al. 2018; Fu et al. 2019). Moreover, based on the framework of metastable systems (Kelso 2012; Tognoli et al. 2014), Naik et al. (2017) suggested the complexity of brain dynamic processes in aging is better understood using the notion of metastability (the ability of the brain to transit between different cognitive states), which could be approximately measured using functional connectivity dynamics. They expected that slow-switching (lower transition probability and/or higher dwell time) in a particular state would be found in the old cohort. A few recent studies began to explore the relationship between dynamic states and age. While it has been discovered that the dwell time of some FC states with weak interactions (Tian et al. 2018) or negative connectivity (Xia et al. 2019) throughout the brain was correlated with age, Tian et al. (2018) did not report the age-related correlation on transition probability. Xia et al. (2019) found age-related increase on transition probability from such states to other states. Chen et al. (2018) only reported that the transition probabilities of specific FC states differed between different age groups, but did not find differences on dwell time. One of the important reasons for the mismatch may be that the aforementioned dFC studies employed the sliding-window (SW) based dFC approaches which suffer from a number of severe limitations such as window length (Hindriks et al. 2016; Leonardi & Van De Ville, 2015), window shape and offset (Shakil et al. 2016), extent of overlap (Betzel et al. 2016), susceptible to sample variability and head motion (Laumann et al. 2016), and lack of reliability (Choe et al. 2017). Moreover, SW-based ‘brain states’ measurements focused on characterizing dynamics of inter-regional correlation, rather than the dynamics of brain activity.. Judging from these concerns, the changes in the metastability of human brain across the adult lifespan may be captured by a more appropriate approach.

Hidden Markov model (HMM) (Rabiner et al. 1989) is one of the promising techniques. HMM models the switching behavior of the brain states as a Markov chain, and assumes that the observed data (i.e. fMRI time series) are generated from these hidden states through a multivariate Gaussian distribution. Therefore, in contrast to the co-activation patterns approach (Liu and Dyun, 2013; Liu et al., 2013; Chen et al., 2015) which directly clustering the single frames of the observation sequence, HMM allows more robust characterization of the dynamic properties.The transition probability between the hidden states, the fractional occupancy (FO) and mean lifetime (the average time spent in each state during each visit) are often used to quantify the temporal dynamics of the inferred hidden states (Baker et al. 2014). HMM has been applied to neuroimaging studies of multiple modalities such as EEG (Hunyadi et al. 2019; Stevner et al. 2019), MEG (Baker et al. 2014; Vidaurre et al. 2016; Quinn et al. 2018; Van Schependom et al. 2019) and fMRI (Chen et al. 2016; Ryali et al. 2016; Vidaurre et al. 2017; Vidaurre et al. 2018; Taghia et al. 2018; Kottaram et al. 2019). Because of the expanded feature space inherent in the probabilistic model, HMM is the most effective when sample size is large due to Chen et al. (2017) and is sufficient to capture quasi-stationary states of activity that are consistently recurring over a population (Baker et al. 2014). The capacity of HMM in discovering the switching dynamics in developmental maturation (Ryali et al. 2016), schizophrenia (Kottaram et al. 2019), multiple sclerosis (Van Schependom et al. 2019) and non-REM sleep (Stevner et al. 2019) proves that HMM is capable of characterizing the dynamic pattern of spontaneous brain activity across the adult lifespan.

In this study, we utilized HMM to estimate subject-specific FO, mean lifetime and transition probabilities of discrete hidden states from rs-fMRI data, and investigated the relationship between these properties and age. Previous studies have revealed the age-difference in the functional interactions between default mode network (DMN) and task-positive networks including cognitive control networks (CCN) (Wu et al. 2011; Geerligs et al. 2015; Keller et al. 2015; Spreng et al. 2018), dorsal attention networks (DAN) (Spreng et al. 2016; Ferreira et al. 2016; Tsvetanov et al. 2016; Esposito et al. 2017; Monteiro et al. 2019), as well as salience networks (SN) / ventral attention networks (VAN) (Ferreira et al. 2016; Tsvetanov et al. 2016; Monteiro et al. 2019). Therefore, we aimed to find significant age-related changes of dynamics within these networks. Moreover, according to the expectation of Naik et al. (2017), we expected to find a slow-switching state with lower transition probability and/or higher dwell time in the old cohort. Furthermore, the temporal clustering approach was included as a control analysis in order to explore the specificity of HMM in uncovering the age effects. Lastly, we aim to find high reproducibility of HMM parameters and their correlation with age.

## 2. Materials and Methods

### 2.1. Participants and image acquisition

#### Dataset 1

The data were selected from Southwest University (SWU) adult lifespan dataset (http://fcon_1000.projects.nitrc.org/indi/retro/sald.html) (Wei et al. 2018). This dataset included 494 healthy subjects, none of which had a history of psychiatric or neurological disorders. The data collection was approved by the Research Ethics Committee of the Brain Imaging Center of Southwest University. The following exclusion criteria were employed: 1) insufficient time points, 2) excessive head motions (max head motion > 2.0 mm or 2.0 degree, or mean framewise displacement (FD) > 0.2 mm) and 3) extremely poor data quality checked by visual inspection. To avoid the uneven distribution of subjects’ ages as well as the imbalance of sex ratio, the left subjects were further divided into six groups according to their ages, with the age of each group spanning nearly 10 years (20–29; 30–39; 40–49; 50–59; 60–69; 70–80). Then we randomly selected 30 subjects, with the constraint that the number of males and females were equal in each group (except for the last group because it only contains 26 subjects (11 males)). In consequence, 176 subjects were included in this study (aged 20-80; mean age = 48.93; SD = 17.03; 86 males).

All of the data were collected at the Brain Imaging Center of Southwest University using a 3.0 T Siemens Trio MRI scanner. Each participant underwent a 3D structural MRI and an rs-fMRI scan. For the functional data, a 8-min scan of 242 contiguous whole-brain images were obtained using a gradient echo echo-planar-imaging (GRE-EPI) sequence with the following parameters: 32 axial slices, repetition time (TR)/echo time (TE) = 2000/30 ms, flip angle = 90 degrees, field of view (FOV) = 220 × 220 mm2, thickness/slice gap = 3/1 mm, resolution matrix = 64 × 64. During the rs-fMRI scan, the subjects were instructed to lie quietly, close their eyes, and rest without thinking about any specific things but to refrain from falling asleep. For the structural data, a magnetization-prepared rapid gradient echo (MPRAGE) sequence was used to acquire high-resolution T1-weighted anatomical images (TR = 1900 ms, TE = 2.52 ms, inversion time (TI) = 900 ms, flip angle = 9 degrees, resolution matrix = 256 × 256, 176 sagittal slices, thickness = 1 mm, FOV = 256 × 256 mm2).

#### Dataset 2

The data were selected from the Enhanced Nathan Kline Institute (NKI) Rockland Sample (http://fcon_1000.projects.nitrc.org/indi/enhanced/index.html) (Nooner et al. 2012) to assess the reproducibility of our findings. This dataset included 941 healthy subjects, none of which had a history of psychiatric or neurological disorders. We only adopted subjects participated in Cross-Sectional Lifespan Connectomics Study. The exclusion criteria were identical with dataset 1. The left subjects were also further divided into six groups according to their ages, with the age of each group spanning nearly 10 years (20–29; 30–39; 40–49; 50–59; 60–69; 70–80). Due to the imbalance of sex ratio, we randomly selected about 30 subjects from each group, and try to include as many male participants as possible. In consequence, 170 subjects were included in this study (aged 20-80; mean age = 49.14; SD = 17.12; 45 males). The age-sex distribution of these 170 subjects could be seen in supplementary Fig. S1.

For the functional data, a 10-min scan of 900 contiguous whole-brain images was obtained using a gradient echo echo-planar-imaging (GRE-EPI) sequence with the following parameters: 40 axial slices, repetition time (TR)/echo time (TE) = 645/30 ms, flip angle = 60 degrees, field of view (FOV) = 222 × 222 mm2, thickness/slice gap = 3/1 mm, resolution matrix = 74 × 74. During the rs-fMRI scan, the subjects were instructed to lie quietly, close their eyes, and rest without thinking about any specific things but to refrain from falling asleep. For the structural data, a magnetization-prepared rapid gradient echo (MPRAGE) sequence was used to acquire high-resolution T1-weighted anatomical images (TR = 1900 ms, TE = 2.52 ms, inversion time (TI) = 900 ms, flip angle = 9 degrees, resolution matrix = 256 × 256, 176 sagittal slices, thickness = 1 mm, FOV = 256 × 256 mm2).

### 2.2. Image preprocessing

The data were preprocessed using data processing assistant for resting-state fMRI (DPARSF, http://rfmri.org/DPARSF) (Yan et al. 2010) with the following steps: removing the first 10 time points, slice timing correction, realignment to the mean functional volume to correct for head motion, spatial normalization to Montreal Neurological Institute space using the T1 image unified segmentation, smoothing (FWHM = 4 mm), nuisance regression including 24 motion parameters (Friston et al., 1996; Yan et al. 2013), linear, quadratic and cubic trends and signals from white matter and cerebrospinal fluid, and bandpass filtering (0.01 - 0.10 Hz).

### 2.3. Definition of functional networks

Our definition of resting-state networks (RSNs) was based on the functional parcellation of the human cerebral cortex with 17 sub-networks (Yeo et al. 2011). The sub-network names were consistent with Betzel et al. (2014) and were shown in Fig.1. For each subject, the averaged time courses within each of the 17 RSNs were extracted from this template and then normalized separately (zero mean and unit standard deviation). The normalized time courses were temporally concatenated across all subjects, resulting in a two-dimensional matrix (networks × (time points × subjects)) to be input into HMM (Fig. 2a).

**Fig. 1.**
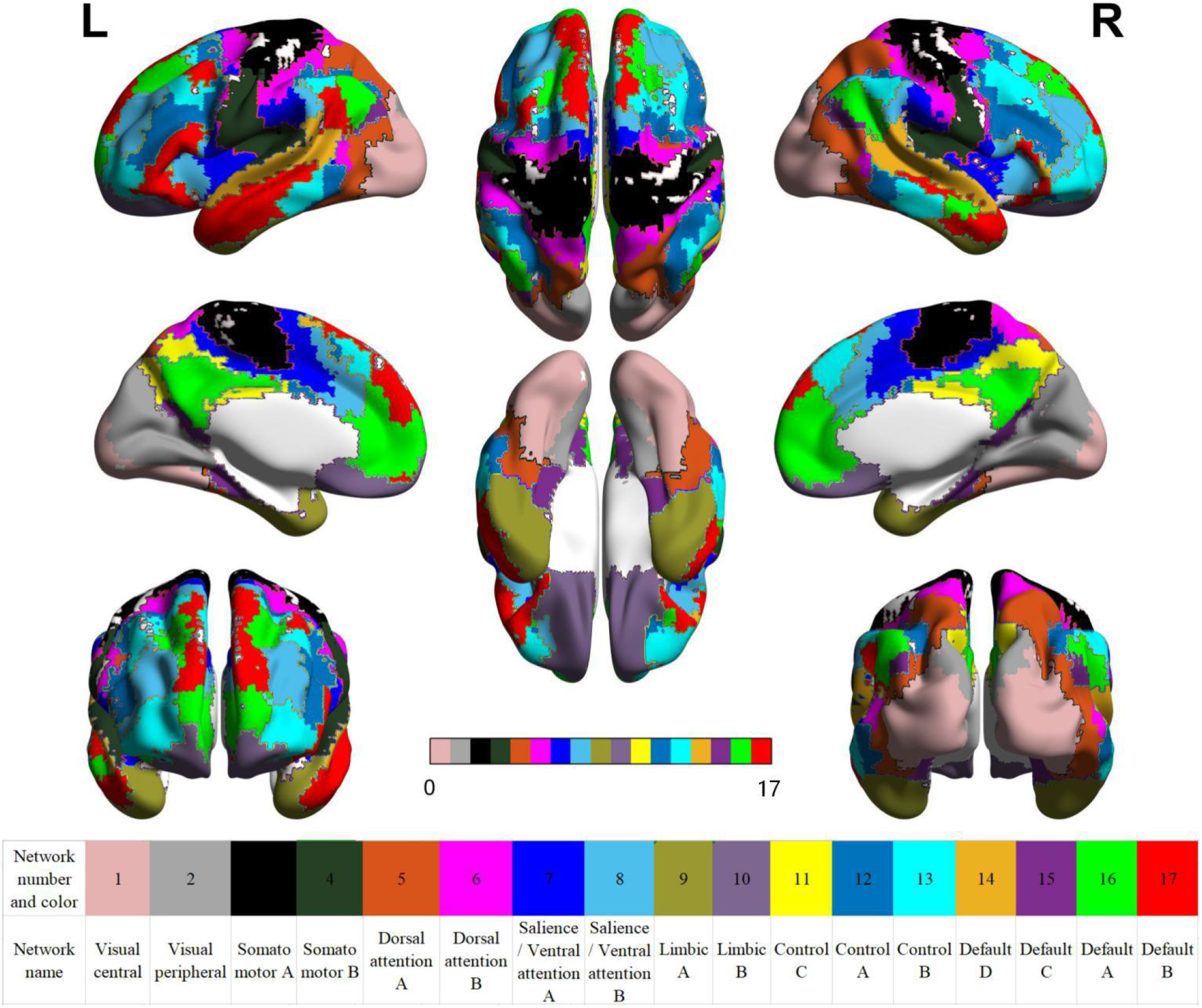
Yeo’s 17-network parcellation of human cerebral cortex. The parcellation are mapped on the smoothed ICBM 152 surface using BrainNet Viewer. Network names are identical with Betzel et al. (2014).

**Fig. 2.**
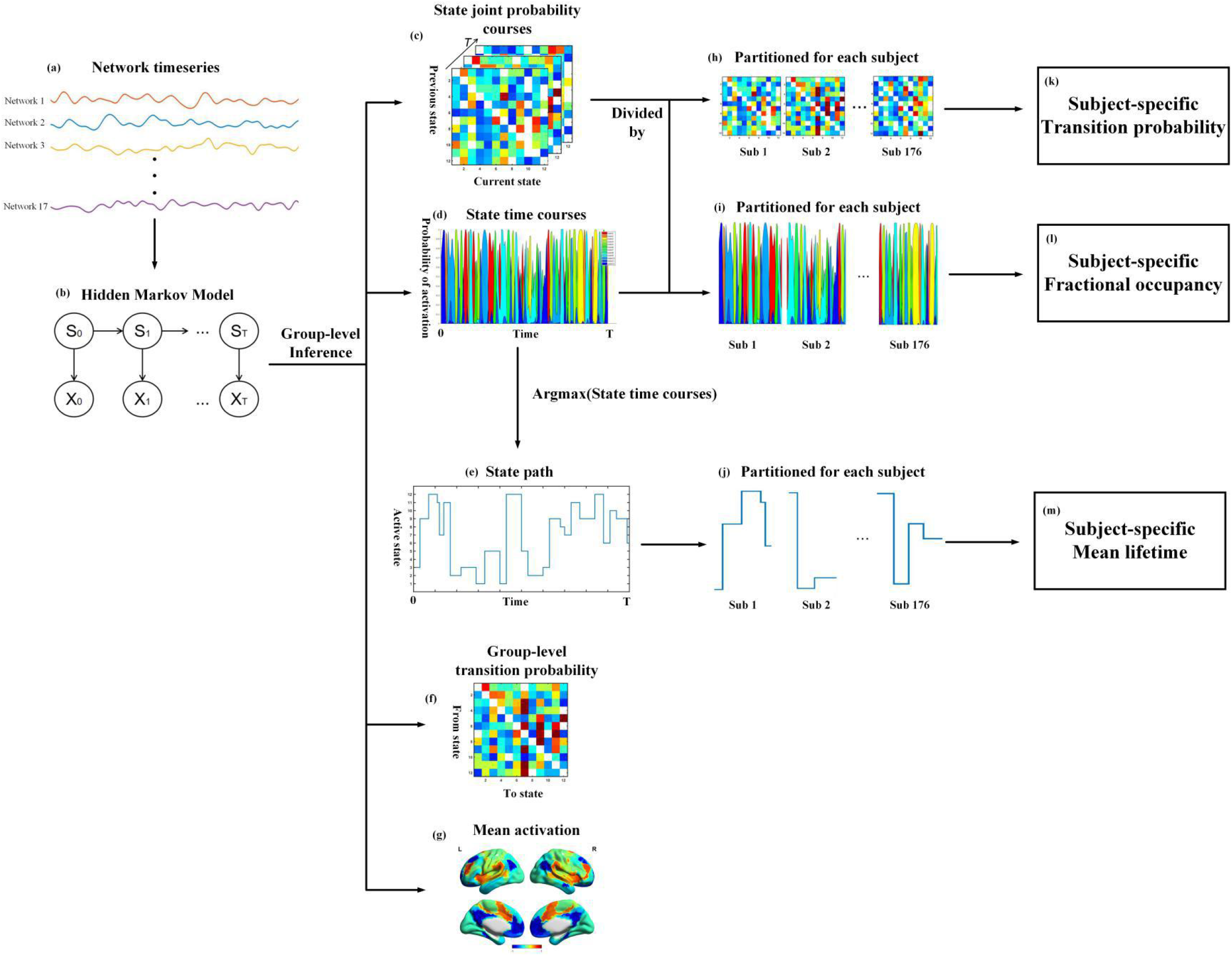
The brief flowchart of our approach. (a) Network-averaged fMRI timeseries were temporally concatenated and fed into HMM. (b) Hidden Markov model. St are random variables following a Markov chain, which denotes the hidden states in time point t. Similarly, Xt denotes the observed values from networks-averaged timeseries in time point t. The conditional probabilities of the observed data on hidden states have the distribution of multivariate Gaussian with mean vectors μ and covariance matrices Σ. (c) State joint probability courses of K = 12 states. Element Aij in each matrix denotes the joint probability P(St,St-1|Xt) of each state conditional on observations at previous time point and at current time point, and is estimated via variational Bayes inference. (d) State time courses of K = 12 states. The posterior probability P(St|Xt)of K states is estimated via variational Bayes inference, with each color representing a particular state. (e) Hidden state path. The state with the maximum posterior probability at each time point. (f) Group-level transition probability matrix. Element Aij denotes the transition probability P(St | St-1) from one state i to another state j. (g) Mean activation of states. Mean vectors μ are projected to the ICBM 152 surface using BrainNet toolbox. (h) The state joint probability courses were divided into subject-specific statistics. (i) The state time courses were divided into subject-specific statistics. (j) The state path was divided into subject-specific statistics. (k) Subject-specific transition probability matrix. Element Aij denotes the transition probability P(St | St-1) from one state i to another state j for each subject. (l) Subject-specific fractional occupancy, which is defined as the temporally averaged posterior probability of state k in one scan. (m) Subject-specific mean lifetime, which is defined as the average amount of time spent in each state before transitioning out of that state for each participant.

### 2.4. Hidden Markov model

A brief flowchart of the HMM approach is shown in Fig. 2. HMM assumes that brain activity (observation sequence), is generated by a number of hidden states (state sequence) that are considered to be random variables following a Markov chain (Fig. 2b). The HMM can be fully determined by a set of basic parameters including 1) initial probability distributions of the hidden states, 2) the transition probability matrix of the hidden states (In a homogeneous Markov chain, probabilities of the current states only depend on those at the previous time point and the transition probability matrix) and 3) the observation model that stands for the conditional probabilities of the observed data on the hidden states and here is assumed to follow multivariate Gaussian distribution with the mean μ and covariance matrix Σ.

In this study, the HMM parameters were estimated by the HMM-MAR toolbox (https://github.com/OHBA-analysis/HMM-MAR) (Vidaurre et al. 2016; Vidaurre et al. 2017) using improved variational Bayes inference (Rezek and Robert, 2005). We ran HMM for 400 cycles with different initialization and chose the cycle with the lowest free energy. Previous HMM studies suggested that 8-12 states were suitable to get physiologically reasonable as well as replicable findings (Vidaurre et al. 2017; Vidaurre et al. 2018; Kottaram et al. 2019). Moreover, for both datasets, the free energy changed negligibly when the number of state was above 12 (Supplementary Fig. S2a-d). Here, the HMM with 12 states was inferred in accordance with Vidaurre et al. 2017 and Kottaram et al. 2019.

By applying the variational Bayes inference on the temporally concatenated data, we firstly estimated the state joint probability courses (i.e., the joint probability of each state at previous time point and at current time point, Fig. 2c), state time courses (i.e., the posterior probability of each state at each time points, Fig. 2d), the state path (i.e., the state with the maximum posterior probability at each time point, Fig. 2e), the group-level transition probability (Fig. 2f), the mean activation (Fig. 2g) and the covariance matrix. The covariance matrix was not analyzed in this study. Next, we divided the state joint probability courses, state time courses and state path into subject-specific statistics (Fig 2h-j). For example, the subject-specific transition probability was calculated by dividing the temporal sum of the state joint probability courses for each participant by the temporal sum of the state time courses of that participant (Fig. 2k) (Rabiner et al. 1989). Subject-specific FO was calculated by temporally averaging the posterior probability of state k for each participant (Fig. 2l), which denotes the proportion of time spent visiting each state (Vidaurre et al. 2017). Subject-specific mean lifetime was defined as the average amount of time spent in each state before transitioning out of that state for each participant (Fig. 2m) (Baker et al. 2014). For simplification, we also considered the subject-specific transition probability matrix as weighted directed graph, and calculated the out-degree and in-degree of each state by summing up each row and column of the subject-specific transition probability matrix. Thus, lower out-degree of a particular state in the elder indicates decreased transition probability from this state to all other states, and vice versa.

### 2.5. Control analysis

Besides HMM, temporal clustering on single frames (Liu and Duyn, 2013; Liu et al., 2013; Chen et al., 2015) has been widely applied to reveal the transient dynamics of resting brain activity. To further clarify whether the age effect was specific for HMM, we additionally performed temporal clustering on the temporally concatenated time series. In particular, the brain states were identified by classifying the frames using k-means ++ algorithm (1000 iterations, 10 replications). To be more comparable with the HMM results, the number of the states for kmeans++ clustering was consistent with that of HMM (K = 12). For both datasets, the within-cluster sum of squared errors (SSE) also changed negligibly when the number of state was above 12 (Supplementary Fig. S2e-h). Similar with HMM, the subject-specific state variables derived from temporal clustering, including dwell time (the proportion of time spent in each state in one scan), mean lifetime and frequency of transition (the frequency of transition from one state to another in one scan)were then calculated by making use of state time courses for each subject.

### 2.6. Multiple linear regression analysis

We employed a general linear model (GLM) to investigate the relationships between subject-specific model parameters (FO, mean lifetime, transition probability (including its out/in-degree) for HMM; dwell time, mean lifetime and frequency of transition for kmeans++ clustering) and age while taking the confounding effects of mean FD, brain volume and sex into consideration. To minimize the effects of possible outliers, we employed robust regression estimation via iteratively reweighted least squares instead of ordinary least squares (OLS) (Holland et al. 1977; Huber 1981; Street et al. 1988). Ramsey’s regression equation specification error test (RESET) (Ramsey 1969) was used to test the presence of specification error in each regression model, and further determine whether the quadratic effect of age should be included as predictors. One-sample t-tests were performed on the regression coefficient of age (if the model is linear) or age2 (if the model is quadratic) (Betzel et al., 2014). The linear or quadratic effect of age was estimated by the t values of the corresponding regressors. Bon-ferroni correction was applied to correct for multiple comparisons. Subsequently, the adjusted FO (or dwell time), mean lifetime, transition probability (or frequency of transition) were obtained by removing contributions of mean FD, brain volume and sex and were further plotted against age.

## 3. Results

### 3.1. Mean activation patterns of the hidden brain states

#### SWU dataset

Mean activation of the 12 hidden states are illustrated in Fig.3a. Generally, there are no one-to-one mapping between these states and the typical RSNs, but these states could be roughly grouped into three categories according to their mean activation levels. 1) Antagonistic states. For example, in state 3, DMN and limbic regions show high-level activation while DAN, SN/VAN and CCN show low-level activation; In state 5, sensorimotor network (SMN) shows high-level activation while SN/VAN and CCN show low-level activation; In state 8, DMN show low-level activation while other networks show high-level activation. 2) Co-activation/Co-deactivation states. States 9 and 7 exhibit the similar patterns that almost all networks show high/low-level activation. 3) Baseline state. State 10 is the baseline state in that all RSNs show moderate-level activation. The names of these states could be seen in Table 1.

**Fig. 3.**
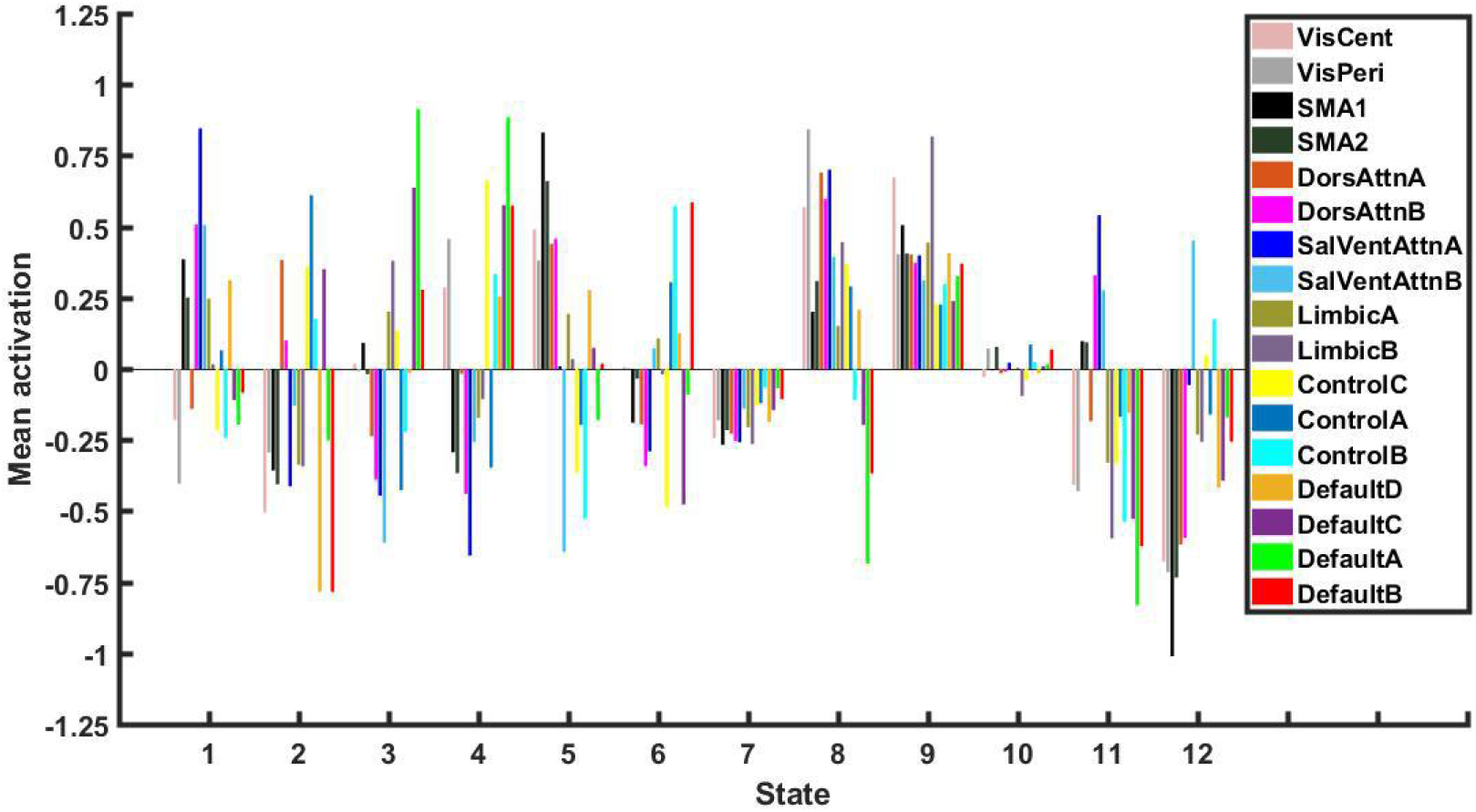
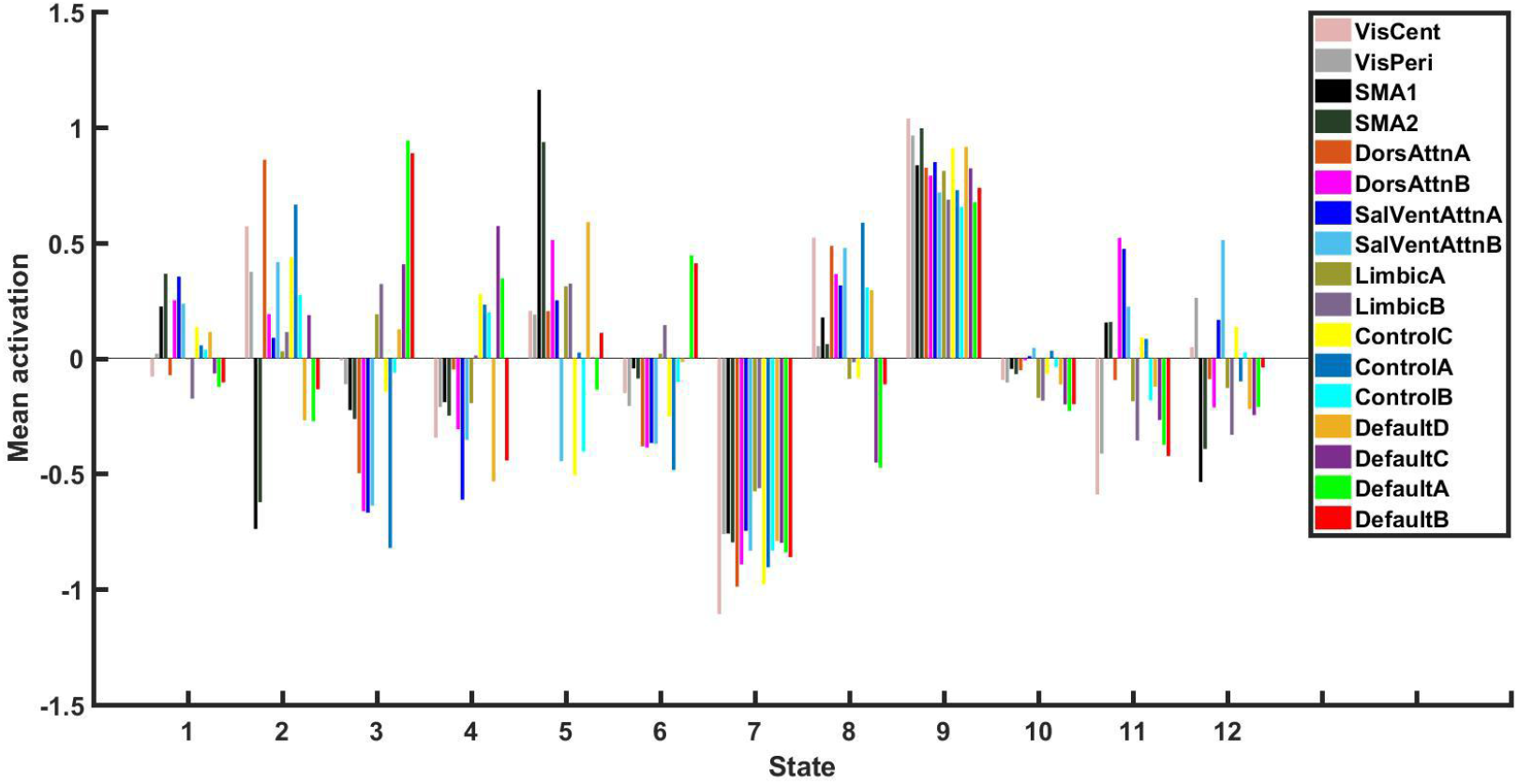
Mean activation of the 12 hidden states in (a) SWU sataset and (b) NKI dataset.

#### NKI dataset

We repeated the same analysis on the NKI dataset. We matched the mean activation maps between the two datasets using Hungarian algorithm based on their distance (L1-norm), and then manually adjusted to avoid mismatch. For dataset 2, mean activation of the 12 hidden states are illustrated in Fig. 3b. It is clear that the relative activation for each state were similar between the two datasets

### 3.2 Temporal parameters of the hidden brain states

#### SWU dataset

Generally, states 7 (All(−)) and 9 (All(+)) have larger FO than the other states (Fig. 4a). However, the averaged mean lifetime are comparable across these states and are about 8 sec (Fig.4b), much less than the typical window length (30-40s) usually used in SW studies. The group-level transition probability matrix, and the averaged out/in degree are illustrated in Fig.4c-e. Notably, FO, mean life time and in degree of state 10 (baseline state) show larger inter-subject variability than the other states. Meanwhile, state 9 (All(+)) and state 10 (baseline state) seem to co-occur with higher head motion than the other state. Interestingly, state 1 (DAN_SN_SMN(+)) and state 10 (baseline state) have larger out degree than the other states.

**Fig. 4.**
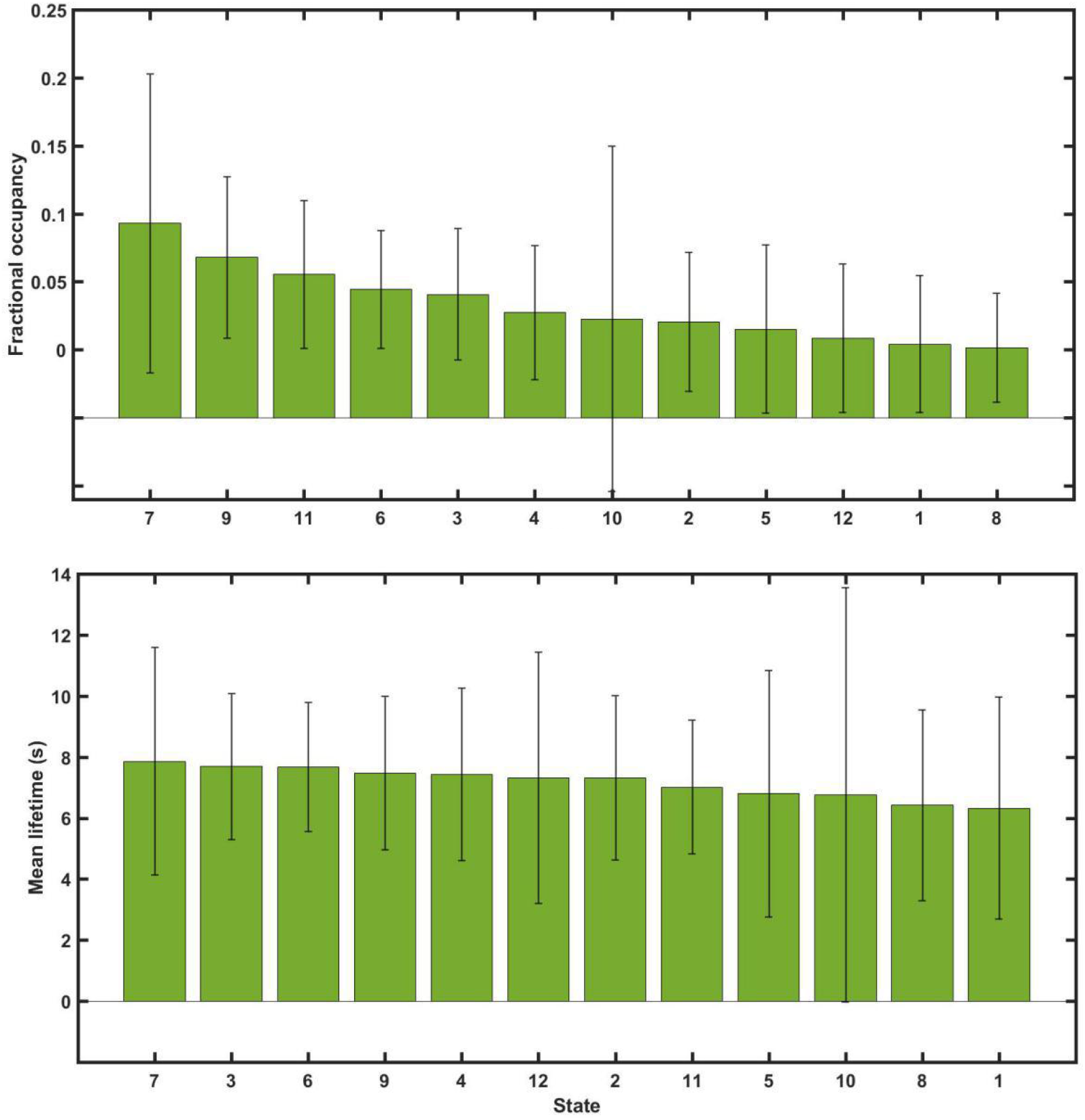
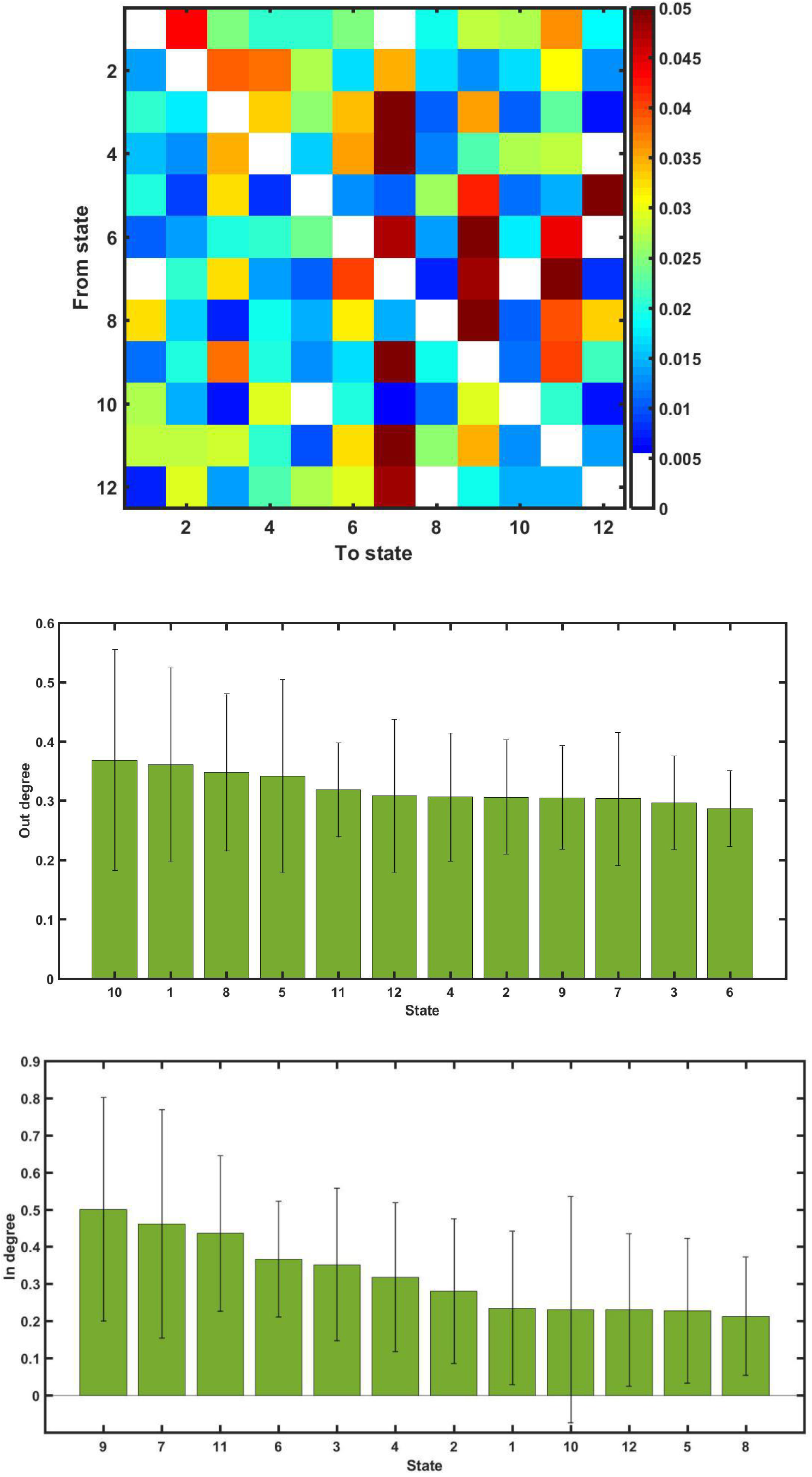
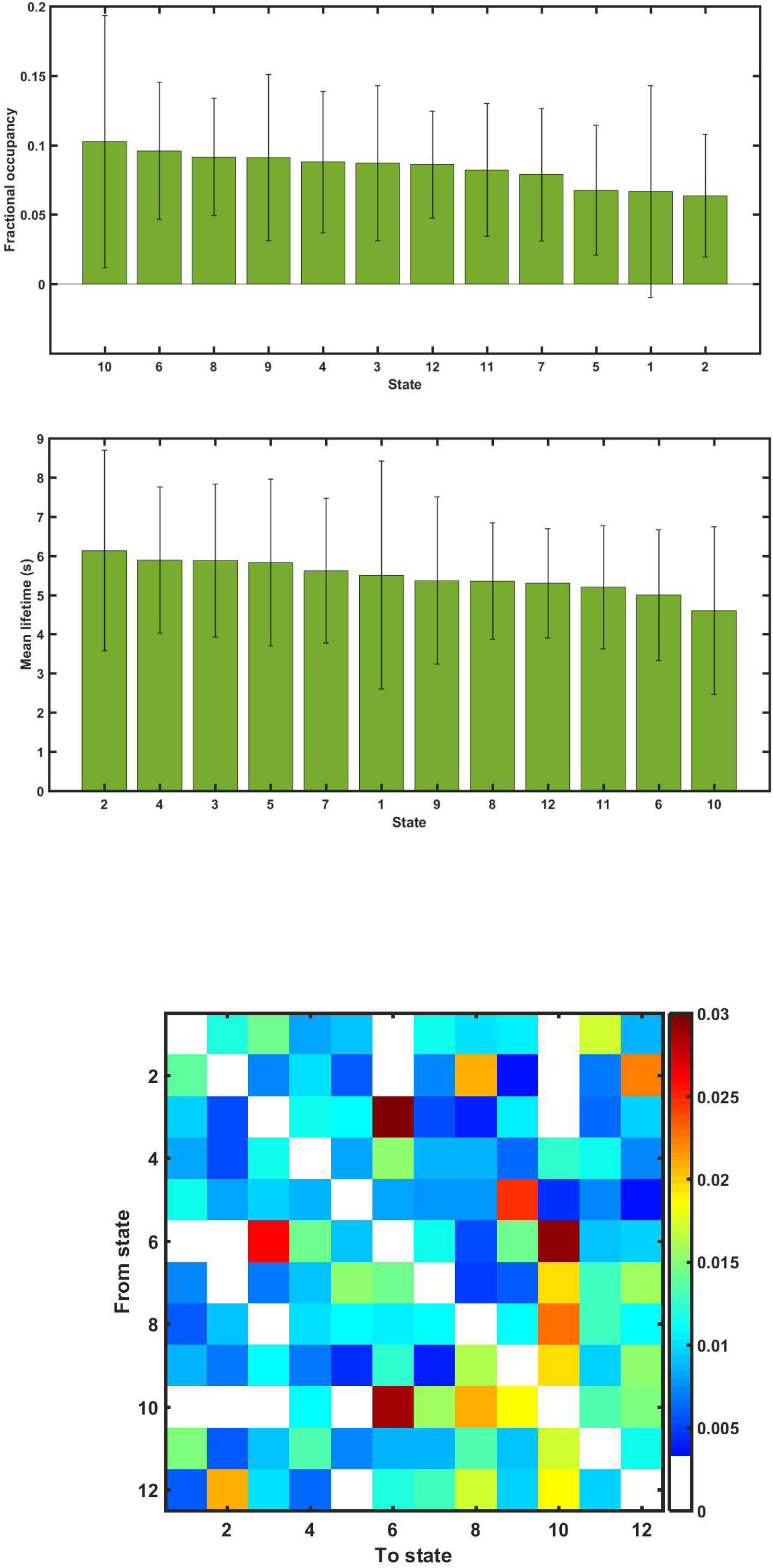
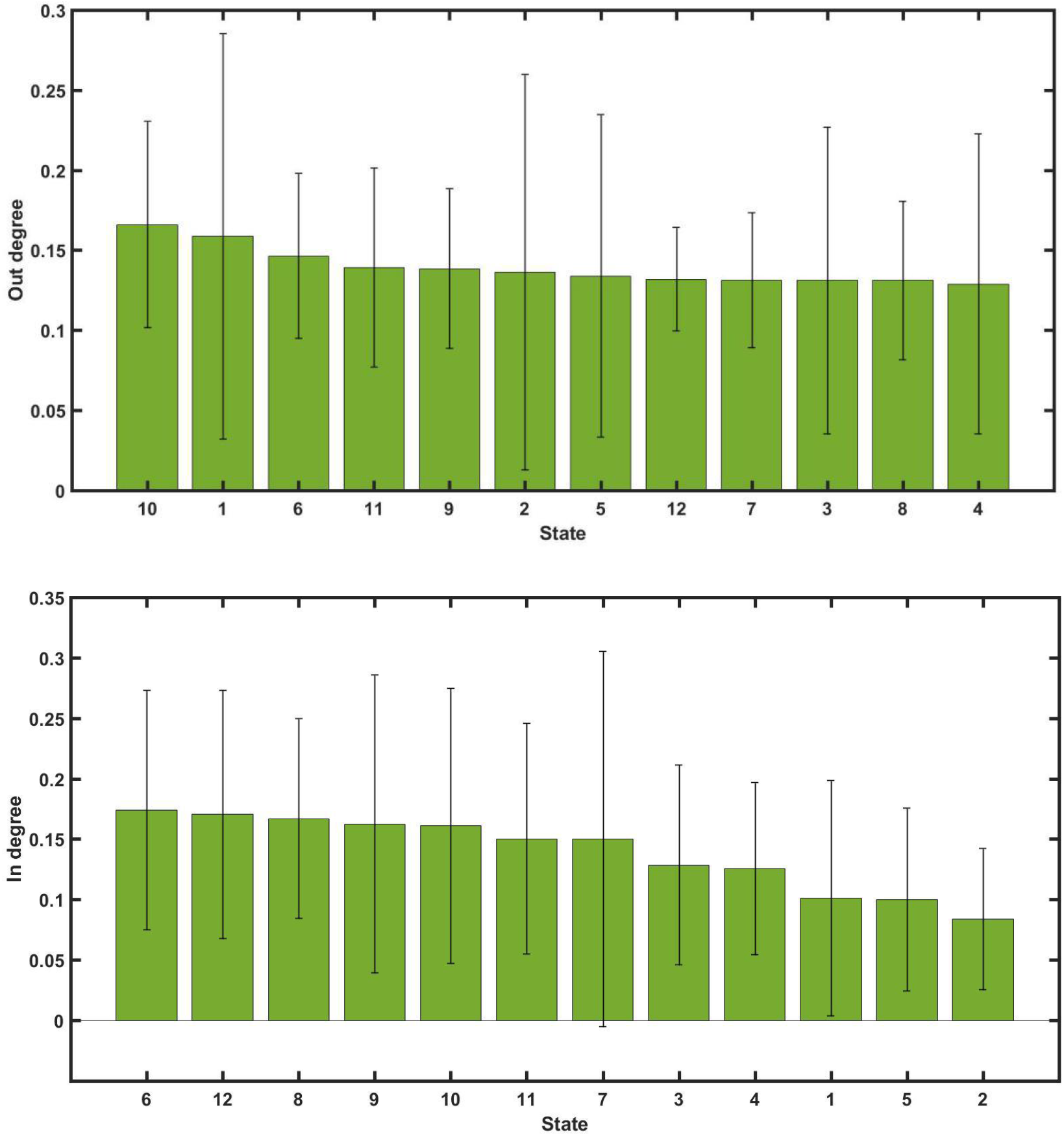
Group-level statistics. (a) Averaged fractional occupancy in SWU dataset. The value of each bar denotes the mean and error bars represent the standard deviation. (b) Averaged mean lifetime in SWU dataset. The value of each bar denotes the mean and error bars represent the standard deviation. (c) Group-level transition probability matrix in SWU dataset. Element Aij denotes the transition probability P(St | St-1) from one state i to another state j. (d) Averaged out-degree in SWU dataset. The value of each bar denotes the mean and error bars represent the standard deviation. (e) Averaged in-degree in SWU dataset. The value of each bar denotes the mean and error bars represent the standard deviation. (f) Averaged fractional occupancy in NKI dataset. The value of each bar denotes the mean and error bars represent the standard deviation. (g) Averaged mean lifetime in NKI dataset. The value of each bar denotes the mean and error bars represent the standard deviation. (h) Group-level transition probability matrix in NKI dataset. Element Aij denotes the transition probability P(St | St-1) from one state i to another state j. (i) Averaged out-degree in NKI dataset. The value of each bar denotes the mean and error bars represent the standard deviation. (j) Averaged out-degree in NKI dataset. The value of each bar denotes the mean and error bars represent the standard deviation.

#### NKI dataset

States 6 (SN_DAN(−)) and 10 (baseline state) have larger FO than the other states (Fig.4f). The averaged mean lifetime are comparable across these states and are about 6 sec (Fig.4g) The FO of state 10 (baseline state) show larger inter-subject variability than the other states. The group-level transition probability matrix, and the averaged out/in degree are illustrated in Fig.4h-j. Meanwhile, state 1 (DAN_SN_SMN(+)) and state 9 (All(+)) seem to co-occur with higher head motion than the other state. Similar with SWU dataset, state 1 (DAN_SN_SMN(+)) and state 10 (baseline state) have larger out degree than the other states.

### 3.2. Age-related changes in FO and mean lifetime of hidden states

#### SWU dataset

The FO of state 3 (DMN(+) / CCN_DAN_SN(−)) exhibits significant and linear decrease with age (Fig.5a, p < 0.05, corrected). The FO of state 10 (baseline state) exhibits significant increase with age and shows a U-shaped lifespan trajectory with the valley around 38 years of age (Fig.5b, p < 0.05, corrected). For the mean life time, only state 10 demonstrates significant increase with age, with a U-shaped lifespan trajectory with the valley around 40 years of age (Fig.5c, p < 0.05, corrected).

**Fig. 5.**
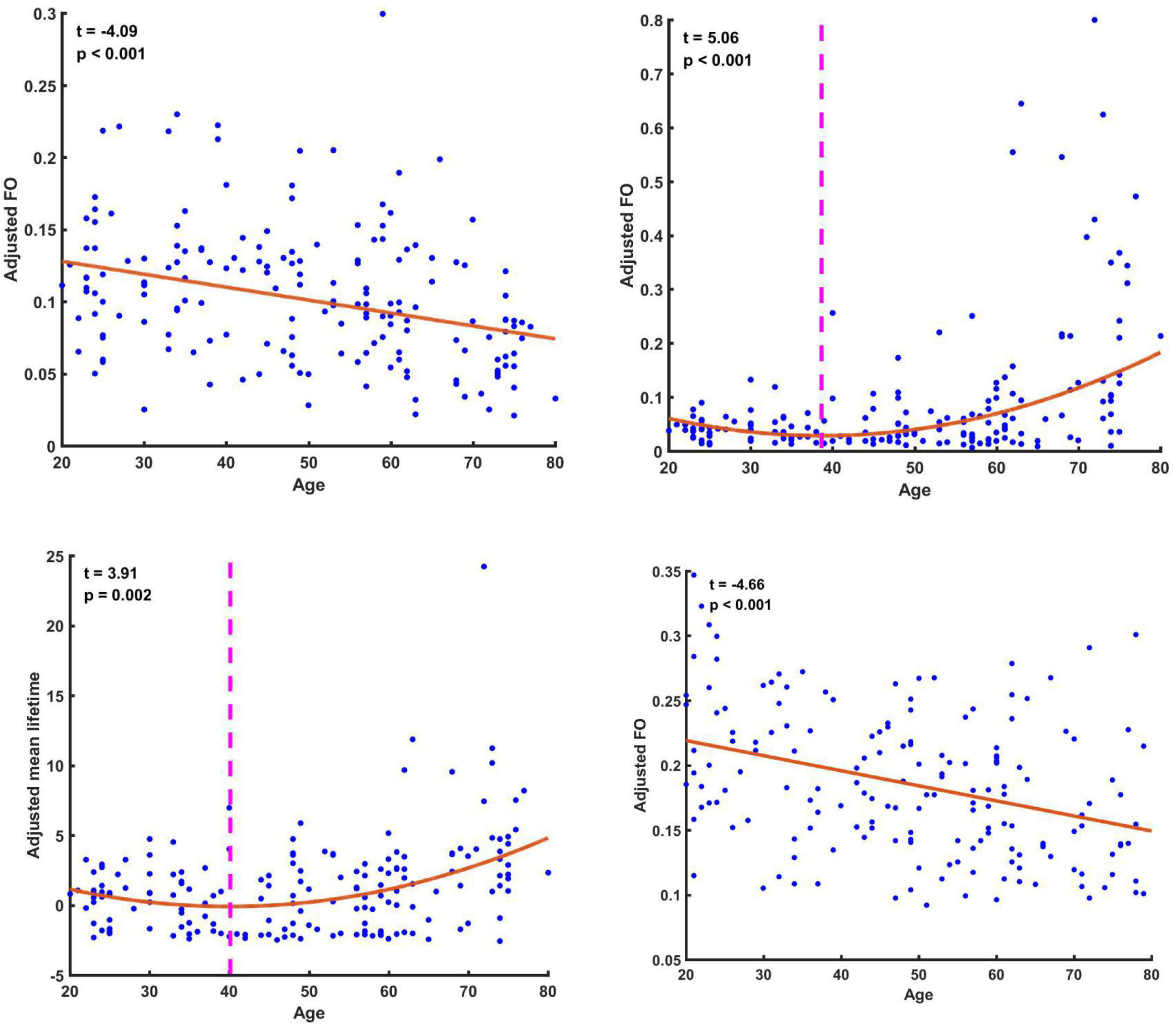
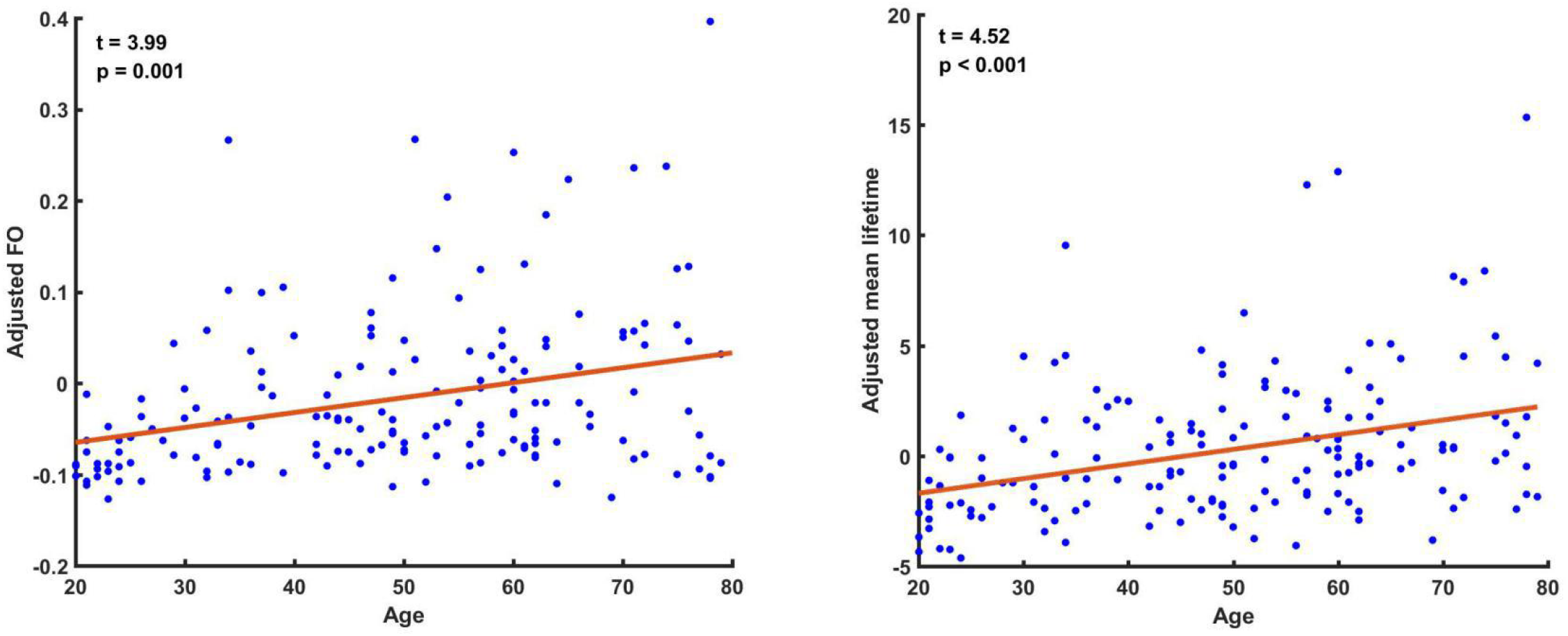
Lifespan trajectories of subjects’ fractional occupancy / mean lifetime. Each dot denotes the adjusted FO / mean lifetime value of each subject. The dashed line denotes the valley of the quadratic fitting curve. (a) Lifespan trajectories of the FO of state 3 in SWU dataset. (b) Lifespan trajectories of the FO of state 10 in SWU dataset. (c) Lifespan trajectories of the mean lifetime of state 10 in SWU dataset. (d) Lifespan trajectories of the FO of state 3 in NKI dataset. (e) Lifespan trajectories of the FO of state 10 in NKI dataset. (f) Lifespan trajectories of the mean lifetime of state 10 in NKI dataset.

#### NKI dataset

We observed similar trends for states 3 (DMN(+) / CCN_DAN_SN(−)) and 10 (baseline state) in the NKI dataset, except that both FO and mean life time of state 10 exhibit linear decrease with age (Fig.5d-f, p < 0.05, corrected).

### 3.3. Age-related changes in transition probability between states

#### SWU dataset

While Out-degree of state 10 (baseline state) in the transition probability matrix decreases linearly with age (Fig.6a, p < 0.05, corrected), the in-degree increases nonlinearly and shows an U-shaped trajectories with age with the peak of 38 years of age (Fig.6b, p < 0.05, corrected). Further analysis reveal that the transition probability from state 2 (CCN_DAN(+) / SMN(−)) and 4 (DAN_SN_SMN(−)) to state 10 (baseline state) increases linearly with age (Fig.6c and d, p < 0.05, corrected). Besides, the transition probabilities from states 9(ALL(+)) to state 10 (baseline state) increase nonlinearly and shows an inverted U-shaped trajectories with age with the peak of 37 years of age (Fig.6e, p < 0.05, corrected). No significant results were observed for the transition probability from state 10 to the other states.

**Fig. 6.**
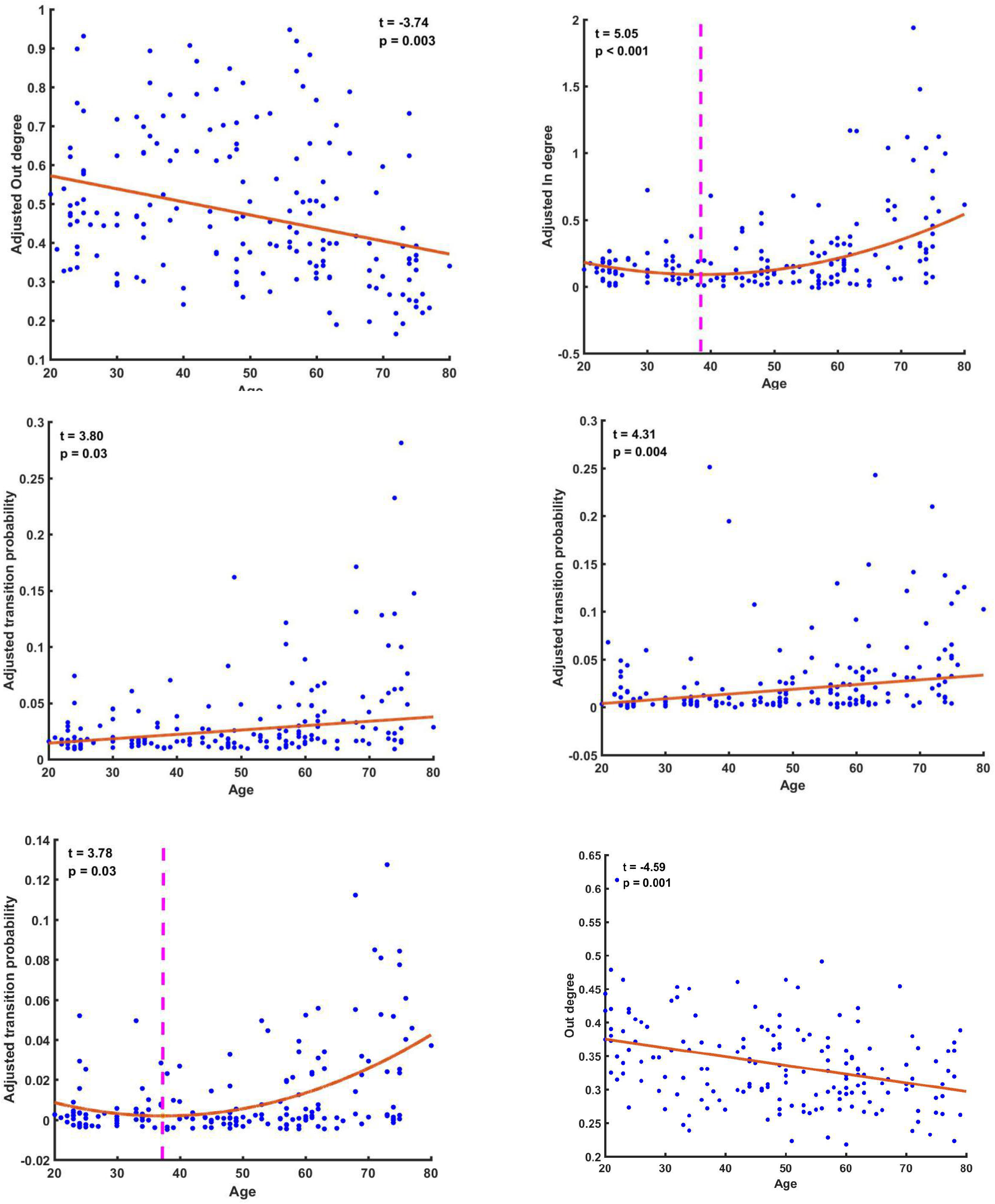
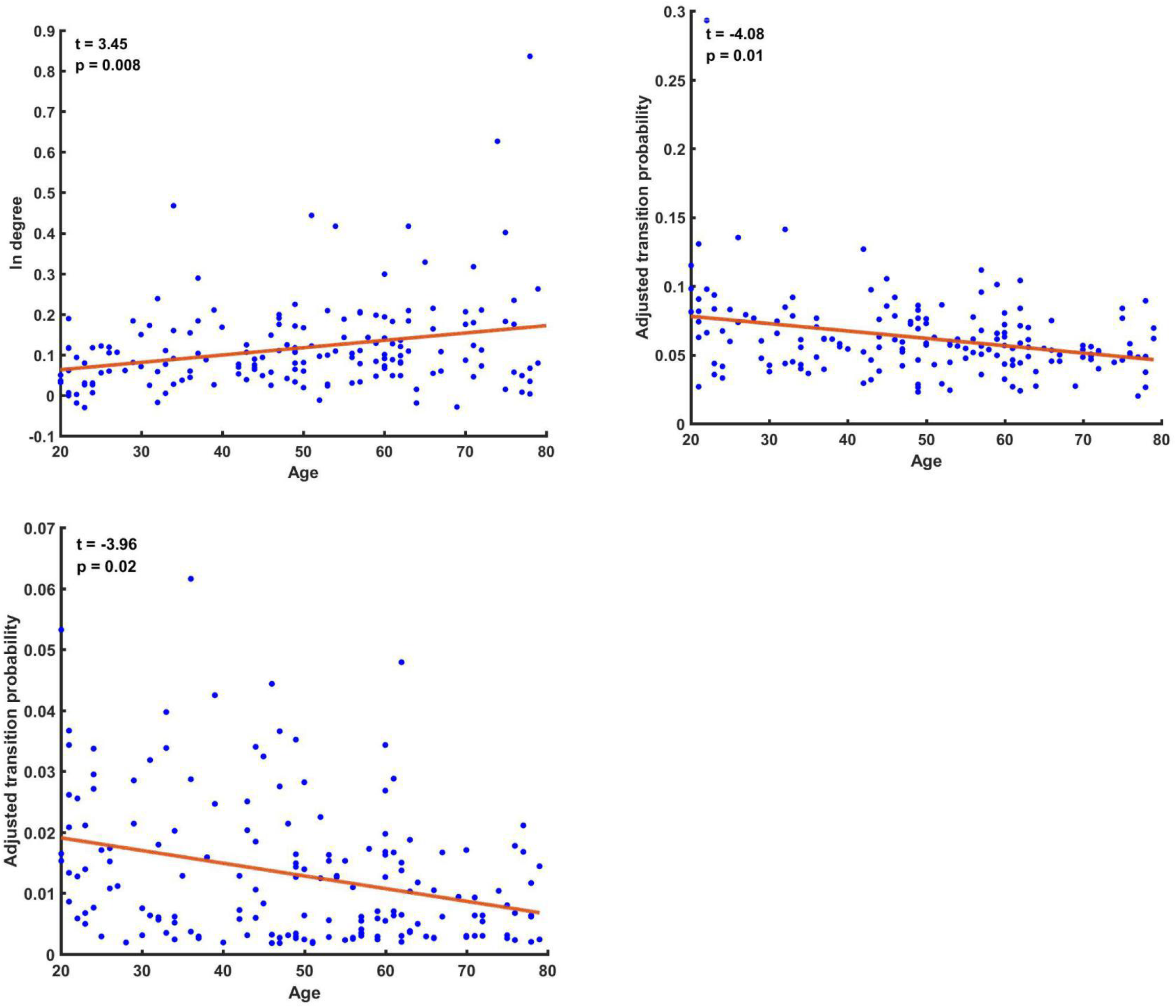
Lifespan trajectories of subjects’ transition probability (and its out/in-degree). Each dot denotes the adjusted transition probability / out/in-degree of each subject. The dashed line denotes the valley of the quadratic fitting curve. (a) Lifespan trajectories of the out-degree of state 10 in SWU dataset. (b) Lifespan trajectories of the in-degree of state 10 in SWU dataset. (c) Lifespan trajectories of the transition probability from state 2 to state 10 in SWU dataset. (d) Lifespan trajectories of the transition probability from state 4 to state 10 in SWU dataset. (e) Lifespan trajectories of the transition probability from state 9 to state 10 in SWU dataset. (f) Lifespan trajectories of the out-degree of state 10 in NKI dataset. (g) Lifespan trajectories of the in-degree of state 10 in NKI dataset. (h) Lifespan trajectories of the transition probability from state 1 to state 6 in NKI dataset. (i) Lifespan trajectories of the transition probability from state 7 to state 8 in NKI dataset.

#### NKI dataset

Out-degree of state 10 (baseline state) in the transition probability matrix decreases linearly with age (Fig. 6f, p < 0.05, corrected), and in-degree increases linearly with age (Fig. 6g, p < 0.05, corrected). Further analysis reveal that the transition probability from state 1 (DAN_SN_SMN(+)) to state 6 (DAN(−)), and from state 7 (ALL(−)) to state 8 (DMN(−) / other(+)) decreases linearly with age (Fig. 6h and i, p < 0.05, corrected)

### 3.4. Control analysis

#### SWU dataset

The kmeans++ results (the spatial maps of the centroids could be seen in supplementary Fig. S3a) show that there is no significance correlation between dwell time/frequency of transition and age. We only found that the mean lifetime of state 3 characterized by deactivation of all networks (Fig. S3a) increases linearly with age (Fig S4a, p < 0.05, corrected)..

#### NKI dataset

The kmeans++ results (the spatial maps of the centroids could be seen in supplementary Fig. S3b) show that there is no significance correlation between frequency of transition and age. We only found that the dwell time and mean lifetime of state 7 characterized by deactivation of all networks (Fig. S3b) increases linearly with age (Fig S4c and d, p < 0.05, corrected) and the dwell time of state 6 characterized by activation of DMN/CCN and deactivation of SN/DAN/SMN (Fig. S4b) decreases linearly with age (Fig S4d, p < 0.05, corrected)

## 4. Discussion

In the present study, we probed into the relationships between the temporal length and transition of discrete brain states, as revealed by HMM, and age using two independent adult-lifespan rs-fMRI datasets. We found significant age-related changes of FO and transition probability in a hidden state involved in higher cognitive and default functions as well as a ‘baseline’ state. The latter corresponded perfectly with the prediction by Naik et al. (2017). Moreover, HMM also exhibited superior specificity in uncovering the age effects when compared with temporal clustering method. Our findings thus suggest that the switching dynamics of spontaneous brain activity may serve as markers of changes in information-processing patterns and higher cognitive functions across the adult lifespan.

### 4.1. Age-correlated decreases in network anti-correlation

Our results showed that the FO of state 3 (DMN(+) / CCN_DAN_SN(−)) decreased with age.. The observation that state 3 (DMN(+) / CCN_DAN_SN(−)) were visited less frequently in the elder suggest that the instances when DMN and task-positive networks show antagonistic behaviors occurs more often in younger than in the elder people. Our results are in agreement with previous static FC studies which reported the reduction of anti-correlation between DMN and CCN/DAN/SN/VAN in the elder (Wu et al. 2011; Geerligs et al. 2015; Ferreira et al. 2016; Keller et al. 2015; Spreng et al. 2016; Tsvetanov et al. 2016; Esposito et al. 2017; Spreng et al. 2018; Monteiro et al. 2019). Evidence suggest that such reduction of anti-correlation was associated with cognitive performance in multiple domains including executive control, processing speed and memory in both younger (Kelly et al. 2008; Hampson et al. 2010; Keller et al. 2015;) and older adults (Putcha et al. 2016; Spreng et al. 2018; Wang et al. 2019). Given that increasing evidences support resting state activity may be primarily determined by brief periods of instantaneous activity instead of a sustained process (Tagliazucchi et al., 2011; Tagliazucchi et al., 2012; Liu and Dyun, 2013; Matsui et al., 2016; Tagliazucchi et al., 2016; Matsui et al., 2019), the reduction of anti-correlation in elder in sFC studies may be primarily attributed to the less discrete brain states as captured by HMM. Of note, there is some conflicts between our findings and two recent studies using SW approaches (Tian et al. 2018; Xia et al. 2019) where the dwell time of states characterized by positive coupling between DMN and CCN (Tian et al. 2018; Xia et al. 2019) and between DMN and SN (Xia et al. 2019) were found to be inversely correlated with age. The discrepancy may be due to that the temporal information of the transient activity is obscured by the SW. For example, the window-length used in Tian et al. 2018 and Xia et al. 2019 are 45 and 44 sec, which are much longer than the averaged mean lifetime of 12 hidden states (about 6-8 sec) in the current study.

### 4.2. Age-correlated changes of ‘baseline’ state

In the current study, we identified a ‘baseline’ state in which all networks only showed moderate-level activity. More interestingly, the FO of this state was positively correlated with age, suggesting that the elder generally spent more time on this state. Such ‘baseline’ state was also found in previous application of HMM into rs-fMRI (Chen et al., 2016; Kottaram et al., 2019). Chen et al. (2016) found that most states often switch to this baseline state and then transit back to other states from it, and thus they named it as ‘ground state’. Different from the findings of Chen et al. (2016), the baseline state observed in the current study was less probable to transit back to other states and tended to occur more frequently in the elder. The discrepancy may be due to the differences in subject ages and scanning parameters between the two studies The subjects used in Chen et al. (2016) were all healthy young adults with an age range of 22–36 years, scanned with a TR of 0.72 s whereas the ages of our subjects span a much wider range and the sampling rate is slower (TR = 2s). While it is possible to limit our analysis to the younger subjects of the lifespan data if we want to control the age effect, the sample size may not be enough for the HMM which requires that the number of time points should be sufficiently large.

The observation that the elder had greater FO in the baseline state than the younger, could be explained by higher transition probabilities from other states to this state, and lower transition probabilities from this state to the others. According to the prediction of Naik et al. (2017), the shift in the dynamic working point (Deco et al. 2013) across the adult lifespan was expected to be reflected as lower transition probability and/or higher dwell-time in a particular network state, which may be due to the age-related changes in structural connectivity (Betzel et al. 2014), synaptic (Morrison et al. 2012) or neurotransmitters (Mora et al. 2007) within some regions, The existence of dynamic working point has been supported by experiment that cortical neuronal networks optimize the ability of information processing when the spontaneous neuronal activity takes the form of ‘neuronal avalanches’(Shew et al. 2009). Thus, decreased transition probability and increased FO of the baseline state in the elder may serve as the compensatory mechanism to ensure neuronal networks optimize the ability of information processing

Of note, the FO of the baseline state demonstrated the U-shaped lifespan trajectory. This suggests the nonlinear relationship between the occurrence of this state and age, as compared to the linear lifespan trajectory of FO for the state DMN_limbic(+) / CCN_DAN_SN(−). Additionally, due to the absence of young adults (age < 30), Tian et al. (2018) found that the dwell time of the loose interaction state increased linearly with age, and expected that the dwell time of the loose interaction state might follow a U-shaped curve throughout lifespan, with children and the elderly spending more time in this state, and the young adults spending less time. And in this study, based on a large number of young adults, we found such U-shaped curve of the occurrence of the baseline state across adult lifespan. Future studies are expected to include sufficient children and adolescents to enrich our knowledge of the developmental trajectories across the whole lifespan.

### 4.3. Specificity of HMM in uncovering the age effects

Comparing with the results of temporal clustering, HMM exhibited higher specificity in uncovering the age effects, especially for age effects in transition probability. This could be explained by the fact that HMM directly models the transition probability, the basic parameter of the Markov chain. By contrast, for the temporal clustering method, state transition is modeled as the frequentist summary statistics of the sequence of brain states. Futhermore, the ‘baseline’ state, which carries distinctive temporal properties, is unable to be captured by methods that only consider spatial similarity, such as k-means used by temporal clustering (Chen et al. 2016). Specifically, we found that the baseline state could not be detected by CAP with different number of states that were frequently used (K = 4 to 15) (Di Perri et al. 2017; Smith et al. 2018; Kaiser et al. 2018; Gutierrez-Barragan et al. 2019; Janes et al. 2020) (Fig. S?). Evidence from previous researches (Chen et al. 2016; Kottaram et al. 2019) and the current study demonstrated that such baseline state could be only detected by using HMM.

From another point of view, our results also suggest that HMM could provide more reliable and biologically plausible results, compared with the SW approaches (Vidaurre et al. 2017; Vidaurre et al. 2018; Kottaram et al. 2019). Specifically, the reliability of SW-based dFC was poor (average ICC < 0.4) and bound to the predefined window size (Choe et al. 2017; Zhang et al. 2018), and was even strongly negatively correlated with the dFC statistical significance (Zhang et al. 2018). Future research is expected to investigate the relationship between structual connections, dynamic working point and the transitions/persistency of HMM states.

Previous studies using sFC have demonstrated the decrease of anti-correlation between DMN and task-positive networks in the elder. While sFC method has merit, it is insufficient to capture the domination and transition of the activation patterns of such anti-correlation between these networks over time, as well as some distinctive, transient states such as the ‘baseline’ state. The current study indicated that HMM is sufficient to capture the anti-correlation patterns across adult lifespan, and further develop the sFC method. However, the results of temporal clustering failed to discover the age effect of the state characterized by the anti-correlation relationship between large-scale networks. On one hand, the results derived from 17 sub-networks show that there is no significance correlation between dwell time/frequency of transition and age, on the other hand, the results derived from 114 ROIs indicate that state 2 and state 12 are not typical ‘anti-correlated’ states due to the low intra-network coupling of DMN/CCN, which largely reduced the interpretability of these states. In brief, compared to sFC/SW/temporal clustering methods, HMM was proved to have incomparable advantages in modelling dynamic properties of adult lifespan due to its direct measurement of state transition and the capability of capturing dynamics of brain activity.

### 4.4 Reproducibility of HMM

Our results demonstrated that similar age effects of the temporal parameters derived from HMM were observed in both datasets. Specifically, the FO of state 3 (DMN(+) / CCN_DAN_SN(−)) and the out-degree of state 10 (baseline state) exhibit significant decrease with age, and the FO, mean lifetime and in-degree of state 10 (baseline state) exhibit significant increase with age, which suggested that changes of the switching dynamics between hidden states underlying observable spontaneous brain activity across the adult lifespan are reproducible and could be potentially widespread. However, compared with HMM, we didn’t find such reproducible age effects in the results derived from temporal clustering, which may be due to the simplicity of k-means method that only considers spatial similarity or defines brain states with frequentist statistics instead of proper probability distributions (i.e. multivariate Gaussian). Future studies are expected to discover more widespread age effects of the switching dynamics of human brain from more numerous and larger datasets.

### 4.5. Limitations

Several limitations should be taken into consideration. Firstly, for both dataset, the mean FD was positively correlated with age (p < 0.05), but we only found significant effect of mean FD on the FO of state 1 (DAN_SN_SMN(+)) and state 9 (ALL(+)) and on the in-degree of state 1 (DAN_SN_SMN(+)) in SWU dataset, and on the FO of state 4 (DAN_SN_SMN(−)) and state 7 (ALL(−)) and on the in-degree of state 4 (DAN_SN_SMN(−)) in NKI dataset in multiple linear regression analysis (p < 0.05, corrected). Besides, state 10 (baseline state) seemed to co-occur with higher head motion than other states in SWU dataset. However, such difference was relatively small and could not be replicated in NKI dataset. Although these issues have been reduced by our strategies of motion-correction, the influence of head motion still remain and could hinder our observations. Secondly, for NKI dataset, the imbalance of sex ratio may induce gender effect. Although we found no significant effect of sex on the FO of 12 states for both datasets, the reproducibility of our study may be limited by the lack of sufficient male participants. Thirdly, we estimated HMM parameters using concatenating data from all individuals, and thus every subject shared a common set of states, which was not suitable to capture subject-specific traits. Future studies are expected to improve the specificity of this approach. Lastly, 400 training cycles of different initialization could not guarantee the global optimum solution, methods like simulated annealing (Lee et al. 2010) or genetic algorithm (Won et al. 2004) may applied to this model and solve this problem.

## 5. Conclusions

This study sharpened our understanding of the link between resting brain dynamics and adult lifespan. Firstly, for two datasets, the time spent on states showing anti-correlation between DMN, CCN and SN/VAN decreased linearly with age. Additionally, for two datasets, we found elder were probably to spend more time on, have less transitions from and have more transitions to an ‘baseline’ state with moderate-level activation of all networks, the time spent on this state also showed an U-shaped lifespan trajectory. Moreover, for both datasets, HMM exhibited higher specificity and reproducibility in uncovering the age effects compared with temporal clustering method, especially for age effects in transition probability. These results demonstrate the age-correlated decrease of the anti-correlation between various networks, and further validate the prediction of Naik et al. (2017) that the existence of a particular network state with lower transition probability and higher fractional occupancy in old cohort, which may reflect the shift of the dynamical working point across the adult lifespan.

## Acknowledgments

This work was supported by the National Natural Science Foundation of China (81201083), the MOE (Ministry of Education in China) Project of Humanities and Social Sciences (16YJCZH057), and the Scientific Research Project of Department of Education of Liaoning Province (LQ2019031).

